# Revealing the benefit of eye motion for acuity under emulated cone loss

**DOI:** 10.64898/2026.03.19.712913

**Authors:** Hannah K. Doyle, James Fong, Ren Ng, Austin Roorda

## Abstract

Retinal degenerative diseases progressively erode the cone photoreceptor mosaic, reducing the retina’s spatial sampling power, yet visual acuity is remarkably resilient to cone loss. Prior work has shown that clinically normal visual acuity (20/25 or better) can persist despite up to 50% of cone cells being lost (Ratnam et al. 2013, Foote et al. 2018). However, studies on individuals with retinal degeneration cannot control for patients’ stage of disease progression, creating the need for an experimental paradigm that can mimic these diseases in healthy subjects. The Oz Vision system creates visual percepts through programmable, per-cell stimulation of thousands of cone cells. We reprogram this system to emulate cone loss in healthy eyes by withholding stimulation from a subset of randomly-selected cones, rendering them inactive, in a method we term “cone dropout.” Using this approach, we characterize the visual system’s robustness to cone loss, showing that visual acuity declines nonlinearly with increasing cone dropout. Importantly, we uncover the compensatory benefit that eye motion provides under cone-deprived conditions, finding that at the highest level of dropout, a visual system with eye motion has an equivalent acuity to a static dropout condition with nearly twice as many sampling elements. Through analysis of eye motion and stimulation data, we find that this benefit arises from the additional information accumulated by “surviving” cones as they sample more of the letter through fixational eye motion.

## Introduction

Retinal degenerative diseases degrade vision by causing photoreceptor death and loss over time (***Georgiou et al., 2021***). Despite this reduction in spatial sampling elements, visual acuity is remarkably robust to cone loss. In patients with retinitis pigmentosa, clinically normal acuity levels (≥20/25) are maintained with up to a 50% reduction in cone density relative to the average (***Ratnam et al., 2013; Foote et al., 2018***). However, study of visual function in patients with retinal degeneration is complicated by the fact that patients may be at different stages of progression in those diseases, the functional status of remaining cones is unknown, and that factors beyond photoreceptor degeneration may contribute to vision loss (***Jones et al., 2016; Lee et al., 2021***). Thus, the mechanism underlying the preservation of acuity under cone loss has not been fully characterized. In light of the challenges posed by studying visual function in patients with retinal degeneration, prior work has attempted to simulate cone loss in healthy subjects. This has historically been achieved through randomly blanking pixels on a display and measuring the impact on subject performance in tasks such as letter identification (***Seiple et al., 1995; Alexander et al., 1995***), grating identification (***Geller et al., 1992; Alexander et al., 1995***), and reading (***Krishnan et al., 2019***). These implementations did not exactly reproduce the conditions of retinal degeneration, though, as the sampling loss was not fixed to the retina as the eye moved and was instead fixed to stimuli in the world. Other studies simulating visual impairment in healthy subjects have accounted for this consideration by stabilizing artificial scotomas to the retina using eye tracking (***Crane and Kelly, 1983; Fine and Rubin, 1999; Kwon et al., 2012; McIlreavy et al., 2012***) and more recently using extended reality technologies in head-mounted displays (***Jones et al., 2020; Chow-Wing-Bom et al., 2020; Sipatchin et al., 2021; Barbieri et al., 2024***). However, these approaches were limited in spatial precision due to tracking error (***Holmqvist and Blignaut, 2020***) and latency between tracking and stimulus updates ranging from 10-80 ms. As a result, the artificial scotomas used in those studies typically spanned multiple degrees of visual angle such that these errors were small by comparison, and thus could not reliably mimic diseases that cause degradation at the much smaller single-cone level.

**Video 1.**
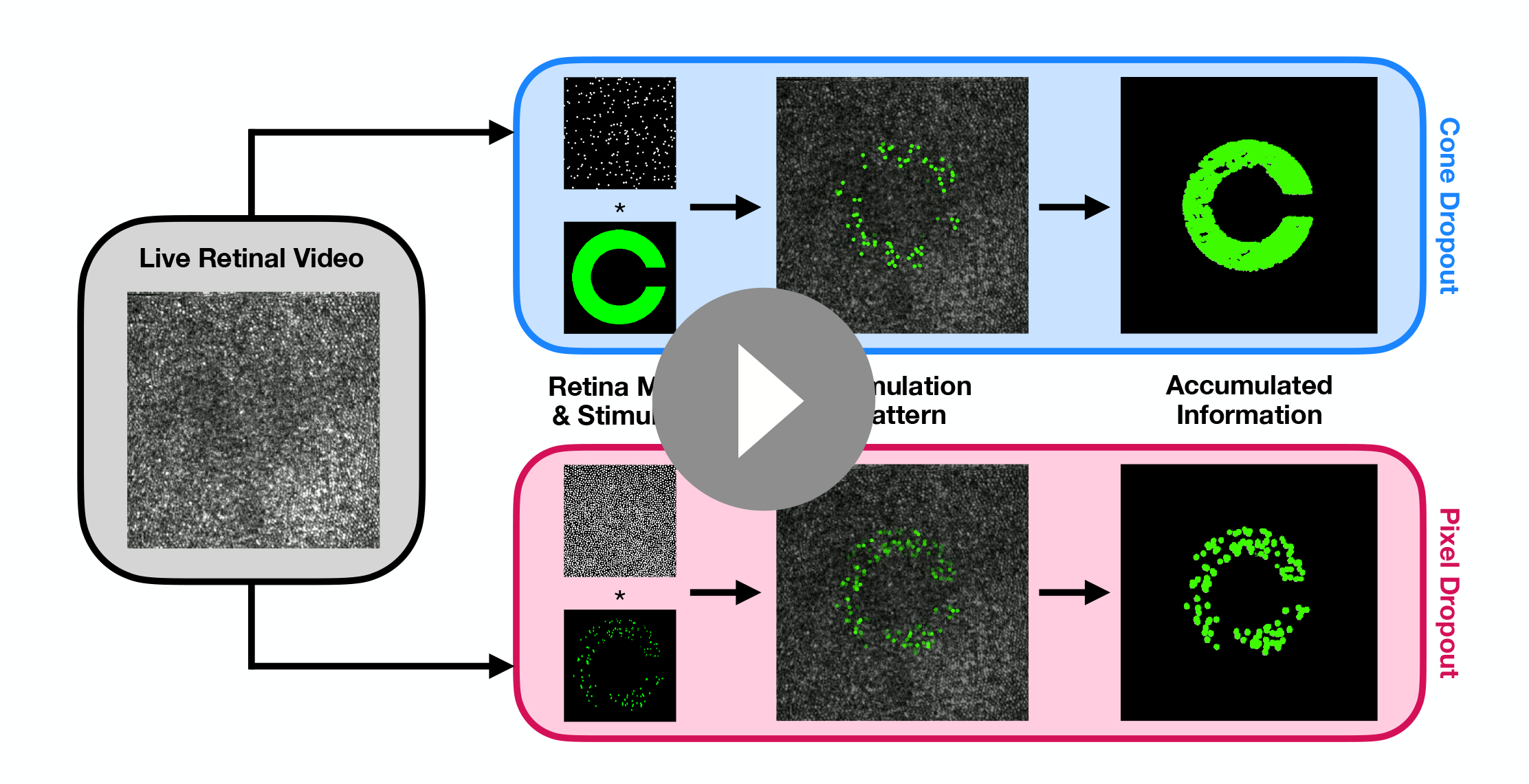
Dynamic comparison between cone and pixel dropout stimulus conditions. The leftmost panel shows a live retinal video from an example trial. The top and bottom panels show visualizations of the cone dropout and pixel dropout conditions respectively. The live retinal image is tracked against a premade retina map where cone locations have been labeled. In the cone dropout condition, a subset of these cone locations have been removed from the map, while the Landolt C stimulus is left unchanged. In the pixel dropout condition, pixels have been removed from the stimulus, while the map remains intact. In each case, the stimulation pattern is the result of sampling the Landolt C stimulus with the underlying retina map as the retinal mosaic moves across the letter. Simulated stimulation patterns are shown for each condition. The final column shows the accumulated information over the course of both example trials. In the cone dropout case, more information is gathered over time due to fixational eye movements, which allow “surviving” cones to sample more of the letter’s area.

The Oz Vision system recreates visual percepts on a per-cone basis through optical stimulation of individual cone photoreceptors at population scales. We implement this principle using an Adaptive Optic Scanning Light Ophthalmoscope (AOSLO), which images the retina, tracks the eye’s motion with sub-cone precision, and stimulates individual cones with 4-ms latency (***Fong et al., 2025***). This technology has previously been used to target cones based on their spectral type, but here we programmatically adapt it to exclude a subset of *all* cones from the stimulation pattern irrespective of their type. That is, we use the system to deliver stimuli cone-by-cone, but delete a subset of randomly-selected cones from an underlying retina map in software such that these cones do not receive stimulation and are rendered effectively “dead.” Then, as the subject moves their eye across the stimulus, only the “surviving” cones are stimulated as they pass under the world-fixed stimulus, and the pattern of “dead” cones remains fixed to their retina. In this way, we can use this system to emulate cone loss that moves *with* the eye, recreating the conditions of retinal degeneration more accurately and on a finer scale than previously possible. While retinal degeneration can induce downstream changes such as retinal rewiring (***Jones et al., 2016; Lee et al., 2021***), in this work, we focus specifically on the impact of cone dropout on visual acuity, and use this stimulation platform to systematically characterize the contribution of photoreceptor sampling alone.

With this technology, we find that vision is adversely affected by cone loss, but that eye motion acts as a compensatory mechanism that preserves visual acuity under a degraded cone mosaic. It has been shown previously that in subjects with normal vision, eye motion can enhance visual acuity (***Ratnam et al., 2017; Rucci et al., 2007; Intoy and Rucci, 2020***). A model developed to explain this phenomenon also predicted that eye motion would provide a benefit for reconstructing a letter E stimulus under conditions of cone loss (***Anderson et al., 2020***). However, no study has measured the effect of eye motion on visual acuity with cone loss in real subjects. We fill this gap by conducting experiments where we programmatically vary the degree of cone loss and measure its impact on visual acuity in a Landolt C task, where subjects must judge the orientation of C stimuli of varying sizes. We compare performance under the condition of emulated cone loss to one where loss is fixed to the stimulus rather than the retina, thereby revealing the role of eye motion in accumulating information when sampling with a degenerated retinal mosaic.

## Results

Subjects performed a Landolt C visual acuity task under two conditions of sampling loss: a “cone dropout” condition, where cones were removed from the retina map, and a “pixel dropout” condition, where pixels were removed from the stimulus itself (see Video 1). In the cone dropout condition, the sampling loss is fixed to the retina and moves with it as the eye moves. In the pixel dropout condition, the loss is fixed to the Landolt C stimulus and does not change over time. The comparison of performance between these two conditions offers insight into the role of eye motion because the percentage of sampling loss present in the stimulus delivered to the retina would be approximately equivalent across the two conditions at any given moment. However, in the cone dropout condition, as the eye moves with fixational eye motion, the visual system would receive information about different regions of the letter C as the intact parts of the retinal mosaic sampled them. Thus, if the visual system makes use of information accumulated over time through eye motion, this would confer a benefit in acuity in the cone dropout condition over the pixel dropout condition.

### Acuity Threshold Experiment

In the first experiment, we measured acuity thresholds using a Landolt C 4-alternative forced choice (4AFC) task. Landolt C stimuli were presented for 1 second, and subjects responded with the orientation of the letter that they perceived (up, down, left, or right). Using randomly-interleaved QUEST staircase procedures to vary the letter size (***Watson and Pelli, 1983***), we determined acuity thresholds under 11 total conditions: 5 percentages of cone dropout, 5 equivalent percentages of pixel dropout, and one no-dropout condition. Each step in these 5 dropout percentages corresponded to reducing the total number of cones or pixels by half.

The results are shown in Figure 1. We plot the measured acuity thresholds for 4 subjects as a function of the fraction of cones remaining in the mosaic and the equivalent percentages of dropout for the cone and pixel dropout conditions. We show the average acuity across all subjects at each percentage tested, and fit linear models to these averages and to each individual subject’s data. Taken as a whole, our data indicates that the cone dropout condition does confer an acuity advantage over the pixel dropout condition spanning approximately one or more rows on a standard acuity chart.

**Figure 1.**
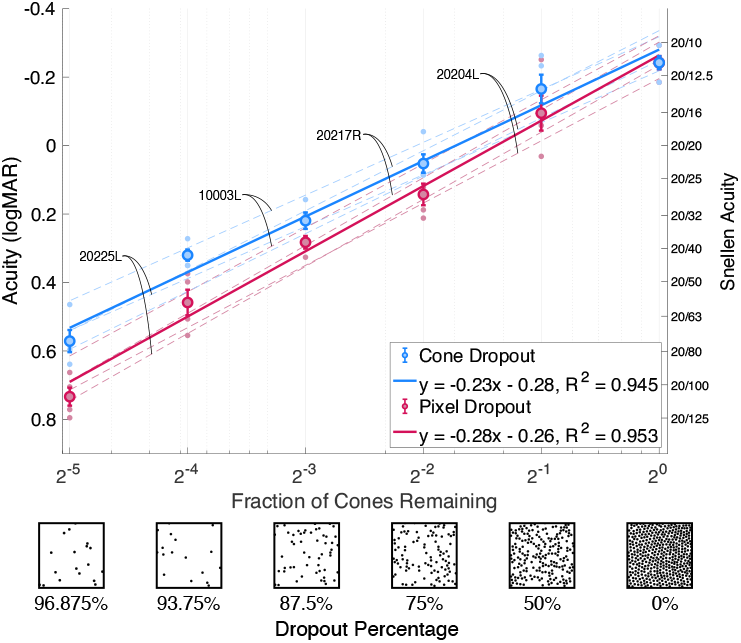
Acuity threshold experiment. Acuity is plotted versus the fraction of cones remaining and the equivalent dropout percentage under the cone dropout condition (blue) and the pixel dropout condition (red) for 4 subjects. Dashed lines show the fitted regression lines to individual subject data, while the solid lines show the fitted regression line across all 4 subjects for the 2 conditions, where *y* is the logMAR acuity and *x* is the logarithm of the fraction of cones remaining in the mosaic. The bold points show the average acuity across subjects with error bars indicating the standard error of the mean.

First, we find that visual acuity in logMAR worsens logarithmically as the dropout percentage increases for the 4 subjects tested (*R*^2^ = 0.945 for cone dropout and *R*^2^ = 0.953 for pixel dropout). This trend is conveyed through linear fits on the log-log plot. The nonlinear relationship between dropout and acuity that we find under emulated cone loss is consistent with prior studies done with real patients, thereby confirming that visual acuity is well-preserved up to 50% cone loss (***Ratnam et al., 2013; Foote et al., 2018***). We use analysis of covariance (ANCOVA) to test for the effects of dropout percentage, dropout type, and the interaction between the two. When looking at the data across all subjects, we find a significant relationship between acuity and dropout percentage (*p* = 3.8 × 10^−30^). There is also a significant difference between acuity on average between the pixel and cone dropout conditions (*p* = 0.00014). Most importantly, the slope of the relationship between acuity and dropout percentage is significantly different between the pixel and cone dropout conditions (*p* = 0.027). That is, eye motion appears to benefit visual acuity when cones are lost from the retinal mosaic.

To contextualize the benefit in acuity provided by eye motion, we indicate the acuity levels corresponding to rows on a standard eye chart on the right side of Figure 1. We can see that at dropout levels of 75% and above, the improvement in acuity from the cone dropout over the pixel dropout condition spans approximately one row or more on the acuity chart. Another feature of the data is that, as dropout increases, the average acuity threshold at a given percentage of cone dropout gets closer to the average acuity threshold at the preceding percentage of pixel dropout. In particular, at the highest level of dropout, the presence of eye motion under cone dropout can effectively recover the acuity level at a pixel dropout condition that has nearly twice as many sampling elements.

### Stimulus Duration Experiment

In a second experiment, we varied the duration of the Landolt C stimulus and measured subject performance for the orientation task under the cone and pixel dropout conditions. We tested 7 durations, spaced logarithmically between 66.7 ms and 4.27 s. The two conditions and 7 durations were randomly interleaved. We kept the dropout percentage fixed at 93.75%, chosen so that the task would be challenging enough to reveal differences in performance across durations and conditions. The letter size was chosen based on each subject’s previously-measured acuity threshold at 93.75% dropout and kept fixed for the entire experiment (see Materials and Methods for details).

Figure 2 shows the results of the experiment varying stimulus duration. We plot Δ Performance versus the stimulus duration, where Δ Performance is the difference in fraction correct between the cone and pixel dropout conditions. This means that points above zero correspond to instances where subjects performed better under the cone dropout condition, whereas points below zero indicate where subjects performed better under the pixel dropout condition. We see that at shorter durations, points are closer to zero, meaning that subjects perform similarly under the pixel and cone dropout conditions. At longer durations, subjects tend toward performing better under the cone dropout condition than in the pixel dropout condition, with the point at the longest duration of 4.27 s being significant (p = 0.018).

**Figure 2.**
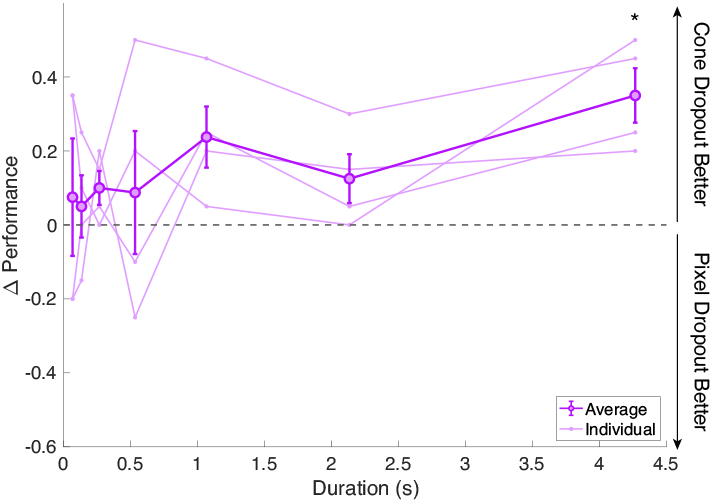
Stimulus duration experiment. The difference in fraction correct between the cone and pixel dropout conditions on a 4AFC Landolt C task is plotted versus stimulus duration.Individual subject Δ Performance is plotted with faint lines. The average Δ Performance across those 4 subjects is plotted with dark lines and error bars indicating the standard error of the mean.

These results provide further insight into the importance of eye motion for recovering information under cone-deprived conditions. At short durations, it makes sense that the two conditions would result in similar performance, as there has been little time for the eye to move and sample new parts of the stimulus under the cone dropout condition. This psychophysical result therefore additionally serves as an experimental validation that the two conditions are perceptually equivalent at a single-frame level. For longer durations,under the pixel dropout condition, no new information could be gained about the stimulus over time. However, the benefit from eye motion in the cone dropout condition was revealed by the performance improvement as stimulus duration increased, as the eye had more time to move an appreciable amount relative to the size of the stimulus.

### Analysis of Recorded Trial Data

In the system we use for cone-by-cone stimulation, a rich stream of data is recorded with every trial, including a video of the moving retina, a record of the eye motion as it was tracked in real time, and a log of all delivered microdoses of light with their corresponding retinal locations. Using this recorded data, we can reconstruct the stimulation pattern that the subject received on any given trial, and conduct analyses in order to help explain our psychophysical results.

#### Sampling Coverage Analysis

In the psychophysical experiments measuring acuity thresholds as a function of dropout percentage, subjects showed better acuity under the cone dropout condition as compared to the pixel dropout condition as dropout increased. The intuitive explanation for such a difference is that under the cone dropout condition, eye motion causes more parts of the letter to be sampled by intact parts of the retinal mosaic, which allows for the visual system to take in more information about the letter, leading to a more accurate judgment of letter orientation. This difference in accumulated information between cone and pixel dropout is depicted visually in Video 1. Using the actual recorded stimulation data, we can examine whether this intuitive understanding is supported by the recorded doses of light that were received by the visual system under the two conditions.

In order to examine the advantage provided by eye motion in the cone dropout condition, we perform an analysis computing the sampling coverage of the Landolt C stimulus on every trial in the experiments measuring acuity thresholds. We define the sampling coverage as the percentage of the letter’s area that was sampled by a cone at any point during a trial (see Materials and Methods for more details on the sampling coverage calculation).

We process every trial delivered during the acuity threshold experiment for all 4 subjects at each dropout percentage under the cone and pixel dropout conditions. We then compute the average sampling coverage across all trials under a given dropout percentage for the two conditions. The results of the analysis are shown in Figure 3a, showing the average sampling coverage versus the fraction of cones remaining. The dashed lines show the fit to each individual subject’s data while the solid lines show the fit across all subjects.

**Figure 3.**
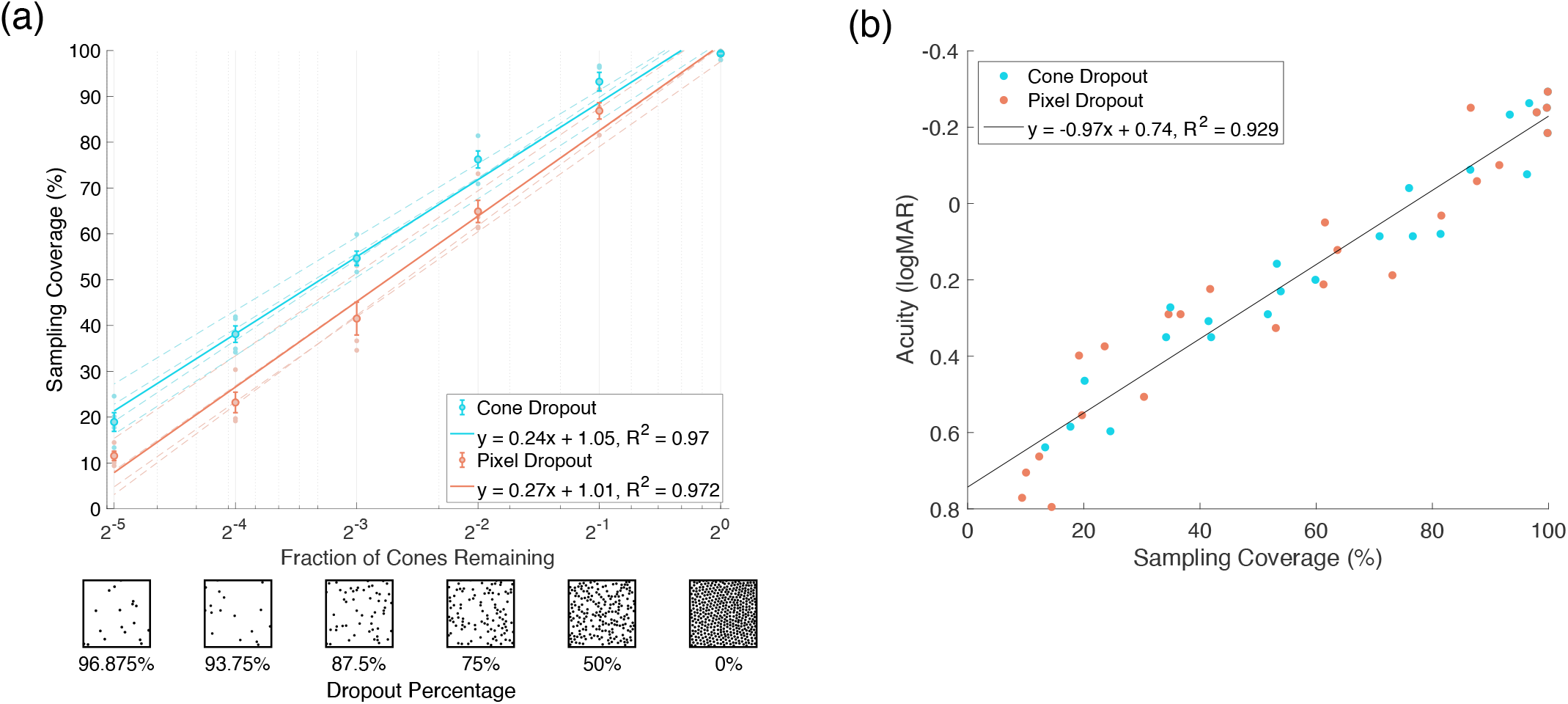
Sampling Coverage Analysis. (a) The average sampling coverage of the Landolt C stimulus across all trials in the acuity threshold experiment is plotted versus the fraction of cones remaining and the equivalent dropout percentage for the cone dropout condition (cyan) and the pixel dropout condition (orange). The dashed lines show the fits to each individual subject’s data, and the solid lines show the fits to the data averaged across subjects, where *y* is the sampling coverage in decimal form and *x* is the logarithm of the fraction of cones remaining in the mosaic. (b) The measured acuity for each subject at each dropout percentage is plotted versus the calculated sampling coverage from the corresponding trials. Each point represents one subject at one dropout percentage, either under the cone dropout (cyan) or pixel dropout (orange) condition. The data is fit by a linear relationship (black), where *y* is the logMAR acuity and *x* is the sampling coverage in decimal form.

We see that the trends in this analysis agree well with the psychophysical results in Figure 1. First, as would be predicted, we see that the sampling coverage is higher under the cone dropout condition (cyan) than under the pixel dropout condition (orange). Additionally, both sets of data are fit well by a logarithmic relationship (*R*^2^ = 0.945 and 0.953 for the psychophysical data and *R*^2^ = 0.97 and 0.972 for the sampling coverage analysis). Qualitatively, the slopes of these best-fit lines appear similar between Figure 1 and Figure 3a. That is, the best-fit line for the pixel dropout condition is steeper than the best-fit line for the cone dropout condition in both cases. These similarities suggest that the quantity sampling coverage, which is calculated directly from stimulation data, can be used as an explanatory metric for the acuities measured during the acuity threshold experiments.

To examine the sampling coverage as an explanatory metric more directly, we plot the measured acuity for each subject under each dropout condition against the corresponding calculated sampling coverage in Figure 3b. We find that the resulting data from both the cone and pixel dropout conditions can be fit well by a single linear relationship (*R*^2^ = 0.929). Importantly, plotting the data in this way causes the acuity gap between cone and pixel dropout to disappear. That is, the difference in measured acuity between the two conditions is captured completely by the difference in sampling coverage. This in turn reflects the key benefit provided by eye motion under the cone dropout condition, where the visual system is able to sample a larger fraction of the occluded Landolt C stimulus.

In this analysis, we sum all microdoses delivered over the course of each 1-second trial, in essence assuming that the temporal integration period of the visual system is infinite. As shown in the experiment varying stimulus duration (see Figure 2), subject performance does seem to improve up to 1 second, after which performance plateaus. Thus, under our experimental conditions, it does seem that the visual system can synthesize all of the additional information gathered over of the course of the 1-second trial. However, in practice, we would not expect for the sampling coverage to continue improving indefinitely with increasing stimulus durations. That is, analysis done for experiments with longer trials should include a finite temporal integration window when computing the sampling coverage.

#### Analysis of Eye Motion Data

The consistent performance gap between the cone and pixel dropout conditions seen in our psychophysical results suggest that eye motion is used to benefit acuity under cone-deprived conditions. Prior work has found changes in eye movement patterns in response to simulated scotomas (***Walsh and Liu, 2014; Yu and Kwon, 2023***) and in patients with retinal degeneration (***Reinhard et al., 2007; Kumar and Chung, 2014***). Thus, we explored the possibility that similar changes could be present in the eye motion traces that were recorded during each experiment.

To determine whether subjects adopted specific gaze strategies in response to the dropout conditions, we examined both the overall magnitude of their eye motion as well as its directional characteristics. To examine the magnitude quantitatively, we computed the iso-density contour areas (ISOAs) containing 68% of the eye motion data from the duration experiment for each subject and dropout condition (see Materials and Methods). Using a two-sample t-test, we found no significant difference in the ISOAs between cone and pixel dropout for all 4 subjects (see Supplementary Figure S1), indicating that subjects did not move their eye any more or less under the cone dropout condition as compared to the pixel dropout condition.

We explore whether directional biases arise in subjects’ eye motion traces using analysis inspired by (***Ratnam et al., 2017***), rotating each eye motion trace as necessary such that all traces are aligned to the veridical orientation of the Landolt C. This aims to reveal whether subjects fixational eye movements tend toward finding the gap in the letter, which they would expect to find at 1 of its 4 possible locations. However, again, we found no evidence of directional differences between dropout conditions or with increasing stimulus duration.

In our analysis of the recorded eye motion traces, we found no clear trends in overall magnitude or directional characteristics, suggesting that the visual system can build a filled-in percept passively through unbiased fixational eye motion. However, it should be noted that in these experiments, the dropout conditions were presented in a random order, meaning that the form of dropout could change between cone and pixel dropout from trial to trial. Thus, it remains to be seen whether a subject would adopt strategic eye motion behavior in response to a fixed pattern of cone dropout that persisted for a longer period of time.

## Discussion

We have developed a technology for faithfully recreating the conditions of cone loss in healthy subjects, and we use it to characterize the relationship between the degree of cone loss and visual acuity. This experimental paradigm solves a set of challenges faced by prior work studying patients with retinal degeneration. Those studies first needed to specifically recruit patients with the diseases of interest. They could not control for the stage of disease progression in each of these patients, and thus were forced to compare patients with a wide range of severity in cone loss. Image-based metrics such as cone spacing may not have been sufficient to know the true degree of cone loss in each patient, as baseline cone density can vary between individuals (***Bensinger et al., 2019***), and even cones that appear dark in a retinal image may still be functional (***Bruce et al., 2015; Tu et al., 2017***). We, however, have complete control over the emulated percentage of cone loss in each of our subjects and can therefore study its effects freely without facing such challenges.

Others have tried to address those challenges by simulating cone loss through randomly blanking pixels in stimuli on a monitor (***Alexander et al., 1995; Geller et al., 1992; Alexander et al., 1995; Krishnan et al., 2019***). One such study even attempted to simulate motion by shifting the blanked pixels midway through the presentation, but lacked the technical capability for stabilization (***Seiple et al., 1995***). Thus, these implementations did not capture the fact that real cone loss is fixed to the retina and will move with the retina due to eye motion. In fact, we have shown that there is indeed a difference in performance between a condition where pixels are removed from the stimulus itself and a condition where cones are removed from the mosaic. That is, using those prior studies as a benchmark would likely lead to an *underestimation* of the degree of cone loss based on a subject’s measured acuity, as those results did not account for the benefit in performance due to eye motion. Therefore, it is critical to consider the impact of eye motion when studying visual function under cone loss.

Our experiments revealed a nonlinear relationship between acuity and the degree of emulated cone loss which resembled the trends found in work with real patients (***Ratnam et al., 2013; Foote et al., 2018***). In particular, our results suggest that one can lose 50% of their cones while maintaining acuity levels of 20/20 or better. This has implications for technologies that aim to restore vision through the use of light-sensitive retinal implants (***Palanker and Goetz, 2018; Holz et al., 2025***). Such technologies consist of a photovoltaic array with an effective sampling density that is much lower than that of a healthy retinal mosaic. The cone dropout condition mimics this loss in sampling power, and thus the acuity levels that we measure can serve as useful benchmarks for the spatial frequency of the array that is required to achieve a target acuity.

In addition to its capability to emulate the conditions of cone loss, our system also captures a stream of data with every trial that can be used to better interpret our psychophysical results. Using this recorded data, we examine both the recorded stimulation patterns and eye motion traces in order to draw conclusions about how acuity improves under the cone dropout condition. We are able to compute what percentage of the Landolt C stimulus was sampled on every trial, and find that this metric best explains the trends in acuity as impacted by dropout condition and percentage. That is, the psychophysical results can be well-predicted directly from the stimulation patterns recorded during each trial.

These experiments mark the first usage of the Oz system for emulating cone loss. In the present study, we apply this technology to characterize visual acuity in the presence of cone loss specifically. However, retinal degeneration has been shown to degrade vision on other metrics such as motion detection (***Turano and Wang, 1992; Alexander et al., 1998***), small-spot detection (***Geller and Sieving, 1993; Makous et al., 2006***), and contrast sensitivity (***Alexander et al., 1992; Lindberg et al., 1981***), and factors beyond the photoreceptors also play a role in shaping vision under retinal degeneration. In this work, we did not model any downstream factors such as shorter outer segments (***Foote et al., 2018***), retinal rewiring (***Jones et al., 2016; Lee et al., 2021***), or ganglion cell hyperactivity (***Kramer, 2023***). Instead, we sought to characterize vision in the presence of cone loss at the lowest possible level, considering only the decrease in sampling power at the retinal input. However, in our system, we can freely vary the experimental task and stimulus conditions to study these aspects of visual function in future work.

In our implementation, we emulate retinal degeneration through the loss of individual cone cells, leaving holes of insensitivity in the retinal mosaic. In reality, the spatial characteristics of photoreceptor loss varies across different retinal degenerative diseases. This can range from contiguous, low density retinal mosaics as in cone-rod dystrophy, simplex retinitis pigmentosa, and neurogenic muscle weakness, ataxia, and retinitis pigmentosa (NARP) syndrome (***Duncan et al., 2007; Yoon et al., 2009***) to patchy, punctate loss as in Stargardt disease, autosomal dominant retinitis pigmentosa, and certain color blindness mutations (***Chen et al., 2011; Duncan et al., 2007; Carroll et al., 2004***). Our approach focuses on punctate dropout and thus does not fully capture the entire range of disease manifestation. However, this technology opens up possibilities for emulating retinal diseases with different forms of dropout. Here, we implement dropout on a cone-by-cone basis, but in general, we have the programmability to fix any pattern of loss to the retina. Future studies could implement intermediate levels of cone degradation rather than binary cone loss, larger spatial scales of dropout, or dropout with varying degrees of spatial uniformity. Thus, this platform enables general study of the way spatial sampling loss impacts visual performance, presenting the opportunity to better understand the implications of a wide range of diseases affecting the retina.

## Methods and Materials

### Subjects

Four experienced male subjects (ages 23-58) were recruited for these experiments, two of whom are authors on this study. All subjects self-reported to have normal vision. Subjects’ peak cone densities varied as follows (subject ID, cone density): 20255L, 11,544 cones/deg^2^; 20217R, 15,748 cones/deg^2^; 10003L, 11,869 cones/deg^2^; 20204L, 11,547 cones/deg^2^. The peak cone density was found using the cone density centroid method described in ***Reiniger et al. (2021***). It involves first generating a cone density map, then computing the density-weighted centroid within the contour at 80% of the peak cone density. All subjects provided informed consent before participating in the experiments. This study was approved by the Institutional Review Board at the University of California, Berkeley.

### Adaptive Optics Scanning Light Ophthalmoscopy

All experiments were run using an AOSLO that simultaneously images and stimulates the retina, while correcting for the imperfect optics of the eye. The adaptive optics component consists of a Shack-Hartmann wavefront sensor which measures the aberrations of the eye, operating in closed loop with a deformable mirror which changes its shape in order to counteract those aberrations. The scanning component of the system consists of two scanners conjugate to the pupil plane, with one resonant scanner operating at 16 kHz in the horizontal dimension and another galvanometric scanner operating at 60 Hz in the vertical dimension. Combined, they scan the outgoing beam of laser light across the retina over a 0.9°-field at a frame rate of 60 Hz, and relay the reflected light back through the system in order to produce a retinal image. The reflected light is detected using photomultiplier tubes.

The AOSLO contains several wavelength channels; in this experiment, we use three of them. Wavefront sensing for the adaptive optics correction was done at 940 nm, retinal imaging was done at 840 nm, and visual stimulation was done at 543 nm.

### Cone-by-Cone Stimulation

In order to deliver cone-by-cone stimulation, we use custom software called Wizard. This software tracks the eye’s motion by breaking the incoming infrared retinal image into strips and locating each strip against a pre-loaded map of the subject’s fovea (more details in *Eye Motion Analysis* section) (***Stevenson and Roorda, SPIE, 2005; Fong et al., 2025***). This map contains metadata that labels the locations of all of the cones within the stimulated region. For the four subjects in this study, the total number of cones in these maps ranged from 17,407 to 26,322. Using these cone locations, Wizard then communicates with an acousto-optic modulator (AOM) to modulate the visible-wavelength channel as it scans across the retina such that it turns on only when it passes over the intended target cones. The latency between the strip used for tracking and the strip where stimulation is delivered in response to that tracking information is approximately 4 ms.

This latency, combined with other factors such as diffraction and residual aberrations, limit the ability to restrict the light to only the targeted cone. Considering a 543-nm focus through a 7.2 mm pupil, a random tracking error with a full-width-at-half maximum (FWHM) of 0.5 arcminutes (***Harmening et al., 2014***), a 0.0125 diopter residual defocus error (maximum error given the step sizes of 0.025 diopters in the AOSLO defocus controller), an average cone spacing of 0.5 arcminutes (***Wang et al., 2019***), and a Gaussian cone acceptance aperture with a FWHM that is 0.5 times the inner segment diameter (***Macleod et al., 1992***), we estimate that each targeted cone receives 5.41 times more light than its nearest neighbor. This means that the ‘dead’ cones cannot be fully excluded from the signal processing. Furthermore, the light leakage reported in ***Fong et al. (2025***) further adds to the signal of non-targeted cones.

Nevertheless, it is important to point out that the information about the stimulus (Landolt C in our case) is sampled at the targeted cone’s location and so, although nearby stimulated cones might detect light, they do not contribute to any increases in the sampling process. This is analogous to adding defocus blur to letters in the pixel dropout condition. Neither situation will improve the spatial information.

### Stimulus Preparation

#### Cone Dropout Condition

In order to implement the cone dropout condition, a randomly-selected subset of the cone labels were removed from the retina map in a proportion equaling the desired dropout percentage. Thus, when the system intended to deliver a C stimulus to the fovea, it would exclude a percentage of the cones from its stimulation as specified by the pre-loaded map.

#### Pixel Dropout Condition

In the pixel dropout condition, we remove pixels from the Landolt C stimulus image. Rather than removing pixels completely randomly, we took measures so that the spatial distribution of the pixel removal would match the spatial characteristics of the cone dropout condition. First, we create a Voronoi tessellation of the subject’s foveal cone map. Next, we randomly select a percentage of the Voronoi cells and create a binary mask, where these chosen cells are set to 0 and the rest are set to 1. We then apply this binary mask to the C stimulus. These stimuli are then delivered through our custom stimulation software, sampled by a retina map containing no dropout. As a result, the pixel dropout stimulus is approximately equivalent to the cone dropout stimulus at any given instant, both in an overall radiometric sense and in terms of its spatial characteristics. This was validated psychophysically by the similarity in performance between the two conditions at short stimulus durations (see Figure 2). The operative difference between the two is that the information loss is fixed to the stimulus in the pixel dropout case, and fixed to the retina in the cone dropout case, which becomes perceptually important as stimulus duration increases.

### Experimental Protocol

#### Experiment Setup

Prior to each experiment, we dilate and cycloplege subjects using drops of 1.0% tropicamide and 2.5% phenylephrine hydrochloride. Subjects are stabilized in the system using a bite bar and moved into alignment using translational stages. For each subject, we determine the best defocus setting for visible stimulation by collecting a series of through-focus images at 543 nm. During each experimental session, we capture a sequence of 0.9° videos in 840 nm to produce a 1.8° × 1.8° map of the fovea which is used for tracking (***Shenoy et al., 2021***). We additionally measure and correct for the transverse chromatic aberration between our imaging and stimulating wavelengths using the method described in (***Fong et al., 2025***).

#### Acuity Threshold Experiment

In the acuity threshold experiment, subjects performed a Landolt C 4AFC orientation identification task. We measured acuity under 5 percentages of cone dropout and 5 equivalent percentages of pixel dropout, along with one no-dropout condition, for a total of 11 conditions. The 5 dropout percentages were logarithmically spaced as follows: 50%, 75%, 87.5%, 93.75%, and 96.875%. For each condition, we ran 4 interleaved QUEST staircase procedures (***Watson and Pelli, 1983***) with 20 trials per staircase, which varied the size of the Landolt C on each trial. We ran equivalent percentages of cone and pixel dropout together, with their staircases randomly interleaved in a block of trials. The no-dropout condition was randomly interleaved with the 50% cone and pixel dropout conditions.

On a given trial, a beep preceded the stimulus, then the stimulus was presented for 1 second, followed by another beep indicating that the subject should respond with their perceived orienta-tion. Another sound played after the subject’s response, and the next trial would begin immediately.

#### Stimulus Duration Experiment

The task in the stimulus duration experiment was the same Landolt C 4AFC task as in the acuity threshold experiment. We tested 7 durations which were logarithmically spaced as follows: 66.7 ms, 133.3 ms, 266.7 ms, 533.3 ms, 1.07 s, 2.13 s, and 4.27 s. These durations correspond to integer multiples of 16.7 ms, the duration of one frame in our 60-Hz frame rate system. Each duration was shown under both the cone and pixel dropout conditions, with 20 trials per duration-dropout combination. All conditions and durations were randomly interleaved.

As in the acuity threshold experiment, a beep preceded the stimulus, then the stimulus was shown for a duration chosen from one of the 7 options, followed by another beep which prompted the subject to a respond. A sound played when the subject’s response was received and the next trial would begin immediately.

In this experiment, we kept the letter size and dropout percentage fixed, and varied only the stimulus duration. We chose the dropout percentage to be 93.75%, so that the dropout was extreme enough to avoid ceiling effects in performance. In order to choose the letter size, we took into consideration all of the data gathered across subjects in the acuity threshold experiment. We normalized each subject’s data to align their psychometric functions by the threshold letter sizes under the cone dropout condition. Then, we fit a single psychometric function to all cone dropout data and to all pixel dropout data separately. We found that the difference in performance between the two conditions was maximized at a 16% larger letter than the threshold. That is, we chose the letter size in the duration experiment to be 16% larger than each individual subject’s previously-determined acuity threshold letter size for the 93.75% dropout condition.

### Eye Motion Analysis

Each trial presented during the experiments has an associated set of data saved along with it. This includes a log of all microdoses that were delivered, the retina map used for tracking, as well as a record of the eye’s motion during the trial.

In the Oz Vision system, each incoming infrared image is broken into 16 strips, which each consist of 16 by 512 pixels. Each strip is located against the premade retina map to produce a record of the eye’s position. There are 16 strips per incoming frame, and our system operates at a frame rate of 60 Hz, resulting in an eye motion trace with a sampling frequency of 960 Hz.

To perform analysis of the eye motion data, we use MATLAB (MathWorks, Natick, MA, USA). With custom software, we iterate through all eye motion data files and extract the x- and y-locations with their associated timepoints. If there are any samples missing due to tracking failures, the missing points are interpolated between the two most recent tracked locations such that there is position data at every timepoint.

With the saved eye motion data, we aggregated all eye motion traces collected during the stimulus duration experiment and separated them by subject and by dropout condition. For each set of eye motion data, we aligned all traces at the centroid of each of their individual distributions. We then computed the iso-density contour that contained 68% of the eye motion data in these aggregated traces, and calculated the area enclosed by that contour. The results of this analysis are shown in Supplementary Figure S1.

### Sampling Coverage Analysis

To compute the sampling coverage for a given trial, we first find the locations of each microdose of light delivered during a given trial. This information is saved in a .csv file with every recorded trial. We create a 2D binary matrix containing a map of each of the pixels where a microdose was delivered. Then, we convolve this map of microdose locations with a 2D Gaussian kernel with *σ* = 0.1 arcmin, in order to simulate the effect of the system point spread function (PSF) which blurs each microdose of light that is delivered to the retina. We binarize the result of this convolution to create a map of all pixels that were sampled over the course of the trial. Finally, we mask this pixel map with a Landolt C with size and orientation corresponding to the current trial, and compute the percentage of pixels covered by the stimulation pattern out of the total possible area contained in the letter. This provides a measure of the sampling coverage of the letter by the subject on each trial.

## Acknowledgments

We thank Pavan Tiruveedhula for technical system support. This research was supported by funding from Air Force Office of Scientific Research grant FA9550-21-1-0230, National Institutes of Health grant R01EY023591 and the Minnie and Roseanna Turner Fund for Impaired Vision Research.

## Notes

### Competing Interest Statement

Ren Ng and Austin Roorda are co-inventors on US Patent Application #17/235627. Austin Roorda is a co-inventor on US Patent #10130253 that is assigned to the University of California.

### Summary of Updates

Minor revisions have been made to the text. A supplemental figure has been added.

## References

Alexander KR, Derlacki DJ, Fishman GA. Contrast thresholds for letter identification in retinitis pigmentosa. Investigative Ophthalmology & Visual Science. 1992 May; 33(6):1846–1852.

Alexander KR, Xie W, Derlacki DJ, Szlyk JP. Effect of spatial sampling on grating resolution and letter identification. Journal of the Optical Society of America A, Optics, Image Science, and Vision. 1995 Sep; 12(9):1825–1833. doi: 10.1364/josaa.12.001825.

Alexander KR, Derlacki DJ, Xie W, Fishman GA, Szlyk JP. Discrimination of spatial displacements by patients with retinitis pigmentosa. Vision Research. 1998 Apr; 38(8):1171–1181. https://www.sciencedirect.com/science/article/pii/S0042698997002356, doi: 10.1016/S0042-6989(97)00235-6.

Anderson AG, Ratnam K, Roorda A, Olshausen BA. High-acuity vision from retinal image motion. Journal of Vision. 2020 Jul; 20(7):34. https://www.ncbi.nlm.nih.gov/pmc/articles/PMC7424138/, doi: 10.1167/jov.20.7.34.

Barbieri M, Albanese GA, Merello A, Crepaldi M, Setti W, Gori M, Canessa A, Sabatini SP, Facchini V, Sandini G. Assessing REALTER simulator: analysis of ocular movements in simulated low-vision conditions with extended reality technology. Frontiers in Bioengineering and Biotechnology. 2024 Apr; 12. https://www.frontiersin.org/journals/bioengineering-and-biotechnology/articles/10.3389/fbioe.2024.1285107/full, doi: 10.3389/fbioe.2024.1285107, publisher: Frontiers.

Bensinger E, Rinella N, Saud A, Loumou P, Ratnam K, Griffin S, Qin J, Porco TC, Roorda A, Duncan JL. Loss of Foveal Cone Structure Precedes Loss of Visual Acuity in Patients With Rod-Cone Degeneration. Investigative Ophthalmology & Visual Science. 2019 Jul; 60(8):3187–3196. https://www.ncbi.nlm.nih.gov/pmc/articles/PMC6657704/, doi: 10.1167/iovs.18-26245.

Bruce KS, Harmening WM, Langston BR, Tuten WS, Roorda A, Sincich LC. Normal Perceptual Sensitivity Arising From Weakly Reflective Cone Photoreceptors. Investigative Ophthalmology & Visual Science. 2015 Jul; 56(8):4431–4438. doi: 10.1167/iovs.15-16547.

Carroll J, Neitz M, Hofer H, Neitz J, Williams DR. Functional photoreceptor loss revealed with adaptive optics: an alternate cause of color blindness. Proceedings of the National Academy of Sciences. 2004; 101(22):8461–8466.

Chen Y, Ratnam K, Sundquist SM, Lujan B, Ayyagari R, Gudiseva VH, Roorda A, Duncan JL. Cone photoreceptor abnormalities correlate with vision loss in patients with Stargardt disease. Investigative Ophthalmology & Visual Science. 2011; 52(6):3281–3292.

Chow-Wing-Bom H, Dekker TM, Jones PR. The worse eye revisited: Evaluating the impact of asymmetric peripheral vision loss on everyday function. Vision Research. 2020 Apr; 169:49–57. https://www.sciencedirect.com/science/article/pii/S0042698920300304, doi: 10.1016/j.visres.2019.10.012.

Crane HD, Kelly DH. Accurate simulation of visual scotomas in normal subjects. Applied Optics. 1983 Jun; 22(12):1802–1806. https://opg.optica.org/ao/abstract.cfm?uri=ao-22-12-1802, doi: 10.1364/AO.22.001802, publisher: Optica Publishing Group.

Duncan JL, Zhang Y, Gandhi J, Nakanishi C, Othman M, Branham KE, Swaroop A, Roorda A. High-resolution imaging with adaptive optics in patients with inherited retinal degeneration. Investigative Ophthalmology & Visual Science. 2007; 48(7):3283–3291.

Fine EM, Rubin GS. Effects of Cataract and Scotoma on Visual Acuity: A Simulation Study. Optometry and Vision Science. 1999 Jul; 76(7):468. https://journals.lww.com/optvissci/abstract/1999/07000/effects_of_cataract_and_scotoma_on_visual_acuity_.22.aspx.

Fong J, Doyle HK, Wang C, Boehm AE, Herbeck SR, Pandiyan VP, Schmidt BP, Tiruveedhula P, Vanston JE, Tuten WS, Sabesan R, Roorda A, Ng R. Novel color via stimulation of individual photoreceptors at population scale. Science Advances. 2025; 11(16):eadu1052. https://www.science.org/doi/abs/10.1126/sciadv.adu1052, doi: 10.1126/sciadv.adu1052.

Foote KG, Loumou P, Griffin S, Qin J, Ratnam K, Porco TC, Roorda A, Duncan JL. Relationship Between Foveal Cone Structure and Visual Acuity Measured With Adaptive Optics Scanning Laser Ophthalmoscopy in Retinal Degeneration. Investigative Ophthalmology & Visual Science. 2018 Jul; 59(8):3385–3393. https://www.ncbi.nlm.nih.gov/pmc/articles/PMC6038831/, doi: 10.1167/iovs.17-23708.

Geller AM, Sieving PA. Assessment of foveal cone photoreceptors in Stargardt’s macular dystrophy using a small dot detection task. Vision Research. 1993 Jul; 33(11):1509–1524. doi: 10.1016/0042-6989(93)90144-l.

Geller AM, Sieving PA, Green DG. Effect on grating identification of sampling with degenerate arrays. Journal of the Optical Society of America A, Optics and Image Science. 1992 Mar; 9(3):472–477. doi: 10.1364/josaa.9.000472.

Georgiou M, Fujinami K, Michaelides M. Inherited retinal diseases: Therapeutics, clinical trials and end points—A review. Clinical & Experimental Ophthalmology. 2021; 49(3):270–288.

Harmening WM, Tuten WS, Roorda A, Sincich LC. Mapping the perceptual grain of the human retina. Journal of Neuroscience. 2014; 34(16):5667–5677.

Holmqvist K, Blignaut P. Small eye movements cannot be reliably measured by video-based P-CR eye-trackers. Behavior Research Methods. 2020 Oct; 52(5):2098–2121. doi: 10.3758/s13428-020-01363-x.

Holz FG, Mer YL, Muqit MMK, Hattenbach LO, Cusumano A, Grisanti S, Kodjikian L, Pileri MA, Matonti F, Souied E, Stanzel BV, Szurman P, Weber M, Bartz-Schmidt KU, Eter N, Delyfer MN, Girmens JF, Overdam KAv, Wolf A, Hornig R, et al. Subretinal Photovoltaic Implant to Restore Vision in Geographic Atrophy Due to AMD. New England Journal of Medicine. 2025; 0(0). https://www.nejm.org/doi/full/10.1056/NEJMoa2501396, doi: 10.1056/NEJMoa2501396.

Intoy J, Rucci M. Finely tuned eye movements enhance visual acuity. Nature Communications. 2020 Feb; 11(1):795. doi: 10.1038/s41467-020-14616-2.

Jones BW, Pfeiffer RL, Ferrell WD, Watt CB, Marmor M, Marc RE. Retinal Remodeling in Human Retinitis Pigmentosa. Experimental Eye Research. 2016 Sep; 150:149–165. https://pmc.ncbi.nlm.nih.gov/articles/PMC5031517/, doi: 10.1016/j.exer.2016.03.018.

Jones PR, Somoskeöy T, Chow-Wing-Bom H, Crabb DP. Seeing other perspectives: evaluating the use of virtual and augmented reality to simulate visual impairments (OpenVisSim). npj Digital Medicine. 2020 Mar; 3(1):32. https://www.nature.com/articles/s41746-020-0242-6, doi: 10.1038/s41746-020-0242-6, publisher: Nature Publishing Group.

Kramer RH. Suppressing retinal remodeling to mitigate vision loss in photoreceptor degenerative disorders. Annual Review of Vision Science. 2023; 9(1):131–153.

Krishnan AK, Queener HM, Stevenson SB, Benoit JS, Bedell HE. Impact of simulated micro-scotomas on reading performance in central and peripheral retina. Experimental Eye Research. 2019 Jun; 183:9–19. doi: 10.1016/j.exer.2018.06.027.

Kumar G, Chung STL. Characteristics of Fixational Eye Movements in People With Macular Disease. Investigative Ophthalmology & Visual Science. 2014 Aug; 55(8):5125–5133. https://doi.org/10.1167/iovs.14-14608, doi: 10.1167/iovs.14-14608.

Kwon M, Ramachandra C, Satgunam P, Mel BW, Peli E, Tjan BS. Contour Enhancement Benefits Older Adults with Simulated Central Field Loss. Optometry and Vision Science. 2012 Sep; 89(9):1374. https://journals.lww.com/optvissci/fulltext/2012/09000/contour_enhancement_benefits_older_adults_with.20.aspx, doi: 10.1097/OPX.0b013e3182678e52.

Lee JY, Care RA, Santina LD, Dunn FA. Impact of Photoreceptor Loss on Retinal Circuitry. Annual Review of Vision Science. 2021; 7(7):105–128.

Lindberg CR, Fishman GA, Anderson RJ, Vasquez V. Contrast sensitivity in retinitis pigmentosa. The British Journal of Ophthalmology. 1981 Dec; 65(12):855–858. https://www.ncbi.nlm.nih.gov/pmc/articles/PMC1039694/.

Macleod DI, Williams DR, Makous W. A visual nonlinearity fed by single cones. Vision Research. 1992; 32(2):347–363.

Makous W, Carroll J, Wolfing JI, Lin J, Christie N, Williams DR. Retinal Microscotomas Revealed with Adaptive-Optics Microflashes. Investigative Ophthalmology & Visual Science. 2006 Sep; 47(9):4160–4167. https://doi.org/10.1167/iovs.05-1195, doi: 10.1167/iovs.05-1195.

McIlreavy L, Fiser J, Bex PJ. Impact of Simulated Central Scotomas on Visual Search in Natural Scenes. Optometry and Vision Science. 2012 Sep; 89(9):1385. https://journals.lww.com/optvissci/fulltext/2012/09000/impact_of_simulated_central_scotomas_on_visual.21.aspx, doi: 10.1097/OPX.0b013e318267a914.

Palanker D, Goetz G. Restoring Sight with Retinal Prostheses. Physics Today. 2018 Jul; 71(7):26–32. https://pmc.ncbi.nlm.nih.gov/articles/PMC6934168/, doi: 10.1063/PT.3.3970.

Ratnam K, Carroll J, Porco TC, Duncan JL, Roorda A. Relationship Between Foveal Cone Structure and Clinical Measures of Visual Function in Patients With Inherited Retinal Degenerations. Investigative Ophthalmology & Visual Science. 2013 Aug; 54(8):5836–5847. https://www.ncbi.nlm.nih.gov/pmc/articles/PMC3757906/, doi: 10.1167/iovs.13-12557.

Ratnam K, Domdei N, Harmening WM, Roorda A. Benefits of retinal image motion at the limits of spatial vision. Journal of Vision. 2017 Jan; 17(1):30. https://www.ncbi.nlm.nih.gov/pmc/articles/PMC5283083/, doi: 10.1167/17.1.30.

Reinhard J, Messias A, Dietz K, MacKeben M, Lakmann R, Scholl HPN, Apfelstedt-Sylla E, Weber BHF, Seeliger MW, Zrenner E, Trauzettel-Klosinski S. Quantifying fixation in patients with Stargardt disease. Vision Research. 2007 Jul; 47(15):2076–2085. https://www.sciencedirect.com/science/article/pii/S0042698907001721, doi: 10.1016/j.visres.2007.04.012.

Reiniger JL, Domdei N, Holz FG, Harmening WM. Human gaze is systematically offset from the center of cone topography. Current Biology. 2021; 31(18):4188–4193.

Rucci M, Iovin R, Poletti M, Santini F. Miniature eye movements enhance fine spatial detail. Nature. 2007 Jun; 447(7146):851–854. doi: 10.1038/nature05866.

Seiple W, Holopigian K, Szlyk JP, Greenstein VC. The effects of random element loss on letter identification: implications for visual acuity loss in patients with retinitis pigmentosa. Vision Research. 1995 Jul; 35(14):2057– 2066. doi: 10.1016/0042-6989(94)00289-x.

Shenoy J, Fong J, Tan J, Roorda A, Ng R. R-SLAM: Optimizing Eye Tracking from Rolling Shutter Video of the Retina. In: 2021 IEEE/CVF International Conference on Computer Vision (ICCV) Montreal, QC, Canada: IEEE; 2021. p. 4832–4841. https://ieeexplore.ieee.org/document/9711255/, doi: 10.1109/ICCV48922.2021.00481.

Sipatchin A, García MG, Wahl S. Target Maintenance in Gaming via Saliency Augmentation: An Early-Stage Scotoma Simulation Study Using Virtual Reality (VR). Applied Sciences. 2021 Aug; 11(15). https://www.mdpi.com/2076-3417/11/15/7164, doi: 10.3390/app11157164.

Stevenson SB, Roorda A. Correcting for miniature eye movements in high resolution scanning laser ophthalmoscopy. In: Ophthalmic Technologies XV, vol. 5688 of Society of Photo-Optical Instrumentation Engineers (SPIE) Conference Series; SPIE, 2005. p. 145–151. doi: 10.1117/12.591190, in.

Tu JH, Foote KG, Lujan BJ, Ratnam K, Qin J, Gorin MB, Cunningham ET, Tuten WS, Duncan JL, Roorda A. Dysflective cones: Visual function and cone reflectivity in long-term follow-up of acute bilateral foveolitis. American Journal of Ophthalmology Case Reports. 2017 Apr; 7:14–19. https://pmc.ncbi.nlm.nih.gov/articles/PMC5644392/, doi: 10.1016/j.ajoc.2017.04.001.

Turano K, Wang X. Motion thresholds in retinitis pigmentosa. Investigative Ophthalmology & Visual Science. 1992 Jul; 33(8):2411–2422.

Walsh DV, Liu L. Adaptation to a simulated central scotoma during visual search training. Vision Research. 2014 Mar; 96:75–86. https://www.sciencedirect.com/science/article/pii/S004269891400008X, doi: 10.1016/j.visres.2014.01.005.

Wang Y, Bensaid N, Tiruveedhula P, Ma J, Ravikumar S, Roorda A. Human foveal cone photoreceptor topography and its dependence on eye length. Elife. 2019; 8:e47148.

Watson AB, Pelli DG. Quest: A Bayesian adaptive psychometric method. Perception & Psychophysics. 1983 Mar; 33(2):113–120. https://doi.org/10.3758/BF03202828, doi: 10.3758/BF03202828.

Yoon MK, Roorda A, Zhang Y, Nakanishi C, Wong LJC, Zhang Q, Gillum L, Green A, Duncan JL. Adaptive optics scanning laser ophthalmoscopy images in a family with the mitochondrial DNA T8993C mutation. Investigative Ophthalmology & Visual Science. 2009; 50(4):1838–1847.

Yu H, Kwon M. Altered Eye Movements During Reading With Simulated Central and Peripheral Visual Field Defects. Investigative Ophthalmology & Visual Science. 2023 Oct; 64(13):21. https://www.ncbi.nlm.nih.gov/pmc/articles/PMC10584020/, doi: 10.1167/iovs.64.13.21.

