## Supplemental Figure. for "Revealing the benefit of eye motion for acuity under emulated cone loss"

Hannah K. Doyle[1], James Fong[1], Ren Ng[1], Austin Roorda[2]
[1]Department of Electrical Engineering & Computer Sciences, University of California, Berkeley, Berkeley, CA
[2]Herbert Wertheim School of Optometry & Vision Science, University of California, Berkeley, Berkeley, CA

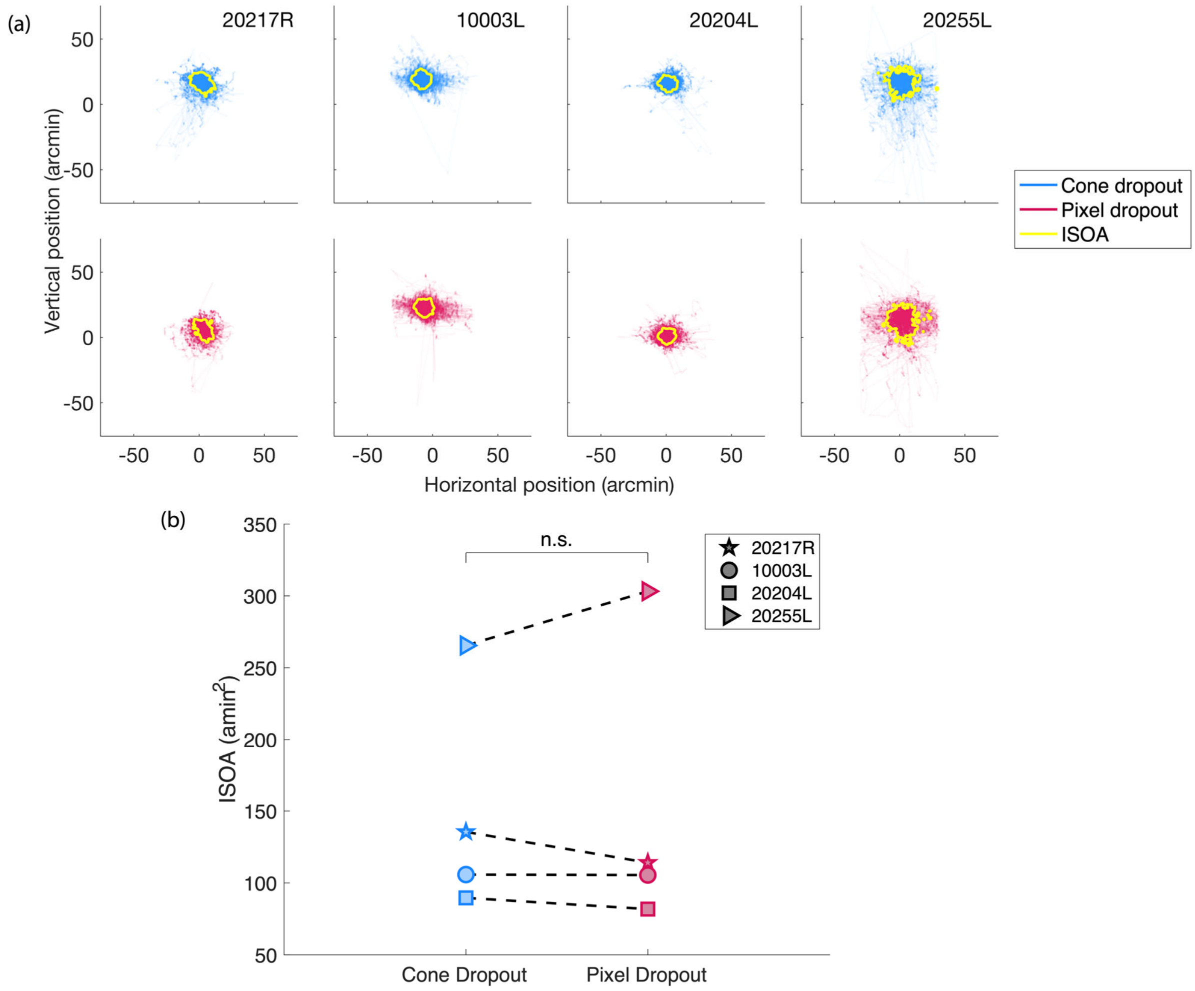


**Figure S1**. Iso-density contour area comparison between dropout conditions. (a) Eye motion traces are plotted for the cone dropout condition (blue) and pixel dropout condition (red) in the experiment varying stimulus duration for 4 subjects as indicated in the upper right corner of each column. The iso-density contour line containing 68% of the eye motion data is plotted on top of the eye motion traces. The apparent cutoff for some traces in the horizontal dimension arises when the subject's eye moves outside of the trackable region. (b) The iso-density contour area (ISOA) enclosed by the contours is plotted for the cone dropout (blue) and pixel dropout (red) conditions. Each symbol indicates a different subject. No significant difference was found between ISOA under the cone dropout and pixel dropout conditions.
